# The zinc-binding motif in tankyrases is required for the structural integrity and proper function of the catalytic domain

**DOI:** 10.1101/2021.10.04.463032

**Authors:** Sven T. Sowa, Lari Lehtiö

## Abstract

Tankyrases are ADP-ribosylating enzymes that regulate many physiological processes in the cell and they are therefore possible drug targets for cancer and fibrotic diseases. The catalytic ADP-ribosyl-transferase domain of tankyrases contains a unique zinc-binding motif of unknown function. Recently, this motif was suggested to be involved in the catalytic activity of tankyrases. In this work, we set out to study the effect of the zinc-binding motif on activity, stability and structure of human tankyrases. We generated mutants of human TNKS1 and TNKS2 abolishing the zinc-binding capabilities and characterized the proteins biochemically and biophysically *in vitro*. We further generated a crystal structure of TNKS2, in which the zinc ion was oxidatively removed. Our work shows that the zinc-binding motif in tankyrases is a crucial structural element which is particularly important for the structural integrity of the acceptor site. While mutation of the motif rendered TNKS1 inactive likely due to introduction of major structural defects, the TNKS2 mutant remained active and displayed a different activity profile compared to the wild type.

## Introduction

Members of the ARTD family are found in all domains of life and are regulators of many processes in the cell such as DNA repair, host-virus interaction, transcription signalling and cellular stress response.^1^ All 17 human ARTD members share an ADP-ribosyltransferase (ART) domain, which is responsible for catalysing the transfer of mono- or poly-ADP-ribosyl groups. Proteins are the most common acceptors of ADP-ribosylation, but also transfer to DNA and RNA was reported for selected family members.^2,3^

Tankyrases (TNKSs) catalyse the transfer of poly-ADP-ribosyl (PAR) groups to their many protein targets.^4–6^ While in lower animals only one TNKS exists, two highly similar TNKSs are present in humans and other vertebrates and are termed tankyrase 1 (TNKS1, PARP5a) and tankyrase 2 (TNKS2, PARP5b).^7,8^ TNKS1 and TNKS2 have overlapping functions in the cell and it is yet unclear what the differences in molecular and physiological functions are.^9,10^ Through their regulatory role in several disease-relevant pathways, TNKSs have emerged as promising drug targets, especially in respect to cancer and fibrotic diseases.^11–13^

In the current model of TNKS function, TNKSs bind their substrate proteins and subsequently poly-ADP-ribosylate (PARylate) them, which leads to PAR-dependent ubiquitination by RNF146 and ultimately proteasomal degradation.^4,14,15^ However, TNKSs also possess non-catalytic functions by acting as scaffolding proteins and mediators of protein-protein interactions, which are under investigation in current TNKS research.^16–19^ Structurally, TNKSs contain five N-terminal ankyrin repeat cluster (ARC) domains. While ARC1-2 and ARC4-5 were shown to be important for binding of their various protein substrates, the ARC3 domain has likely a structural role.^20,21^ A unique feature of TNKSs over other PARP proteins is the presence of a sterile alpha motif (SAM) domain, which allows formation of large multimeric TNKS complexes.^17,22–25^ The ART domain of TNKSs is located at the C-terminus and catalyses the PARylation of substrate proteins.^4,26^

When the first crystal structure of human TNKS1 was determined, it surprisingly revealed a zinc-binding motif present in the catalytic domain located about 20 Å from the NAD^+^-binding pocket.^26^ Presence of zinc in the structure of human TNKS2 was experimentally confirmed by X-ray crystallography shortly thereafter.^27^ Sequence analysis shows that this motif is highly conserved among TNKSs (**Figure 1a**). The zinc-binding CHCC-type motif is located within a loop region of 12 consecutive residues in the ART domain of TNKS1 and TNKS2 (**Figure 1b, c**) and does not structurally resemble known zinc-finger motifs, however the fold can be classified as short zinc-binding loop^28^.

**Figure 1:**
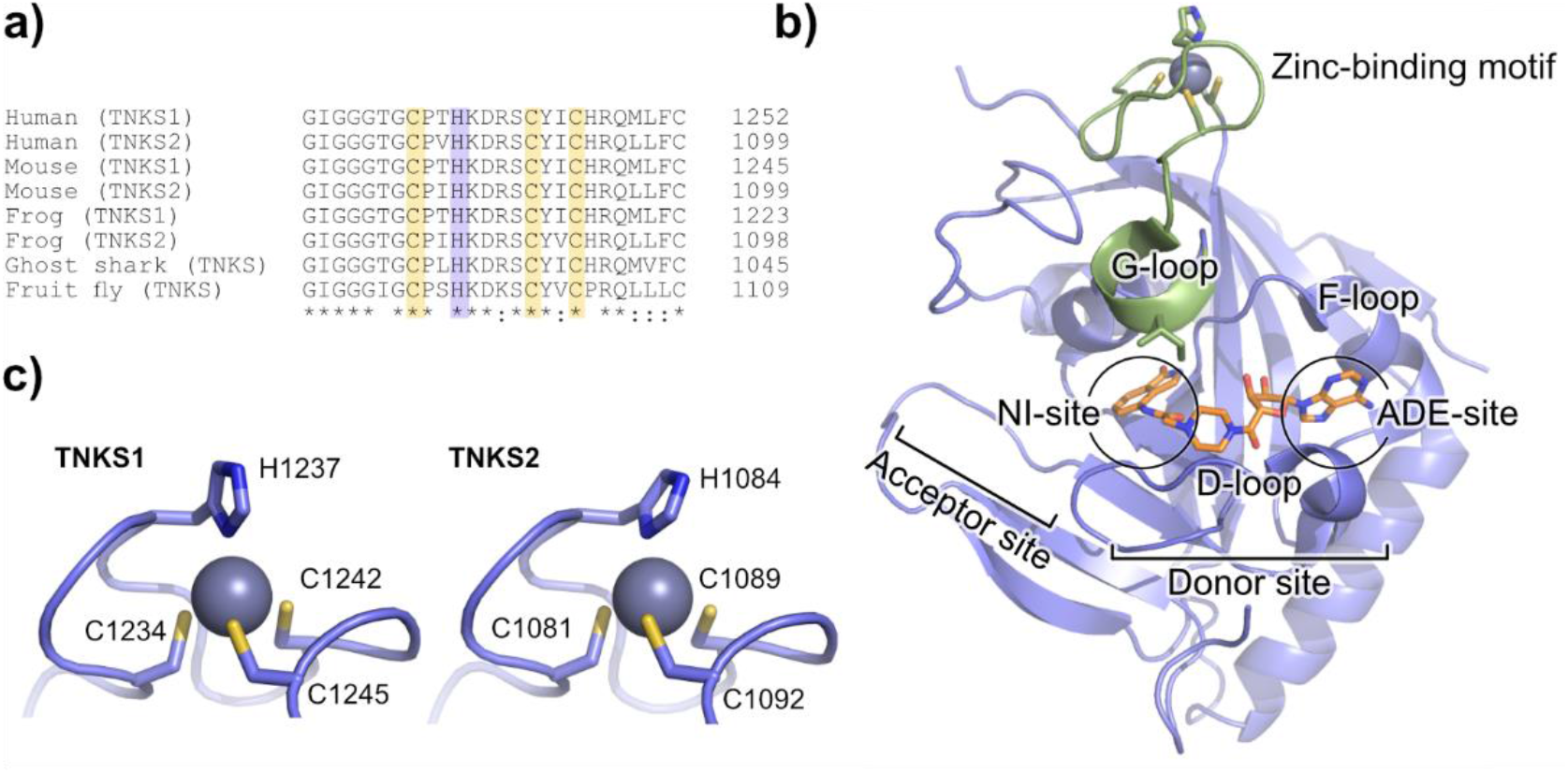
The zinc-binding motif of tankyrases. (a) Sequence alignment of tankyrases. (b) Structure of TNKS2 ART domain containing the zinc-binding motif (PDB: 4BJ9^31^). The zinc-binding motif is structurally connected to the G-loop (green) located close to the acceptor site. The NAD^+^-mimic EB-47 is bound to the active site and is shown in orange. Nicotinamide sub-pocket = NI-site, adenosine sub-pocket = ADE-site. (c) Close-up view of the zinc-binding motif of TNKS1 and TNKS2 with numbered coordinating residues.

No other ARTD family member is known to harbour a zinc-binding function in the catalytic domain, which raised questions about its function in TNKSs. While it was speculated that it could have a structural function, be involved regulation of catalytic activity or protein partner binding^26,29^, no experimental data was available to give more insight into the function. More recently, a study by Kang et al. (2017) found that the motif in TNKS1 can be subject to oxidative damage in the cell resulting in removal of the zinc ion and complete loss of TNKS1 activity.^30^ A direct involvement of the zinc-binding motif in the catalytic mechanism was assumed based on these findings, however it was not discussed how exactly it would contribute to the catalysis based on the TNKS1 structure.

We were intrigued by these results, as the zinc-binding motif is distantly located from the active site. It is however directly connected to the G-loop, which is positioned at the nicotinamide sub-pocket at the acceptor site of TNKSs (**Figure 1b**). While no residues involved in catalysis are located on the G-loop, structurally this loop may still be of importance for NAD^+^ binding and TNKS activity, in which case the zinc-binding motif might provide supporting structural role for the positioning of the G-loop.

In this work, we set out to investigate the function of the zinc-binding motif of TNKSs using biochemical, biophysical and X-ray crystallographic studies. We generated mutants in the TNKS1 and TNKS2 catalytic domains to abolish the zinc-chelating function of the respective motifs and conducted *in vitro* tests on activity and stability. We also generated a zinc-free TNKS2 catalytic domain crystal structure by oxidation of the zinc-binding motif in TNKS2 crystals, revealing structural changes occurring in the TNKS2 ART domain after removal of the zinc ion. Our results show that the zinc-binding motif is an important structural feature in TNKSs and that it is not directly required for the catalytic PARylation activity of TNKSs. However, the motif may support structures such as the G-loop region and can therefore be involved in the control and regulation of the catalytic activity.

## Results

### Activity analysis

We generated and recombinantly produced mutants of TNKS1 and TNKS2 to abolish the zinc-binding capabilities of the motif. In these mutants, the first cysteine and the histidine of the zinc-binding motif were mutated to alanine residues. The mutated residues correspond to Cys1234 and His1237 in TNKS1 and Cys1081 and His1084 in TNKS2 (**Figure 1c**). TNKS activity was shown to be manyfold higher when the SAM-domain present in the construct^17,23^, so we tested the activities of constructs comprising the SAM and ART domains of human TNKS1 and TNKS2. All tests were done *in vitro* with recombinantly produced and purified proteins. For this, we incubated the wild type and mutant constructs of TNKS1_SAM-ART_ and TNKS2_SAM-ART_ with biotinylated NAD^+^ analogs and detected auto-modification of the TNKS constructs by western blot with streptavidin coupled to HRP (**Figure 2a**). The PARylation appears as a high molecular weight smear in western blots. Similar to the report by Kang et al., the mutation in TNKS1_SAM-ART_ completely abolished auto-modification levels compared to the wild type^30^. Interestingly, the mutated TNKS2_SAM-ART_ retained auto-modification activity, however the smear seems to be visible only in lower molecular weight compared to the wild type, possibly indicating formation of shorter PAR chains or modification of fewer acceptor sites. We also visualized total levels of PARylation which might have occurred during expression in *E. coli* by detection with PAR-binder ALC1 fused to nanoluciferase.^32^ Similarly here PARylation smears are detected for all constructs except for the TNKS1 mutant.

**Figure 2:**
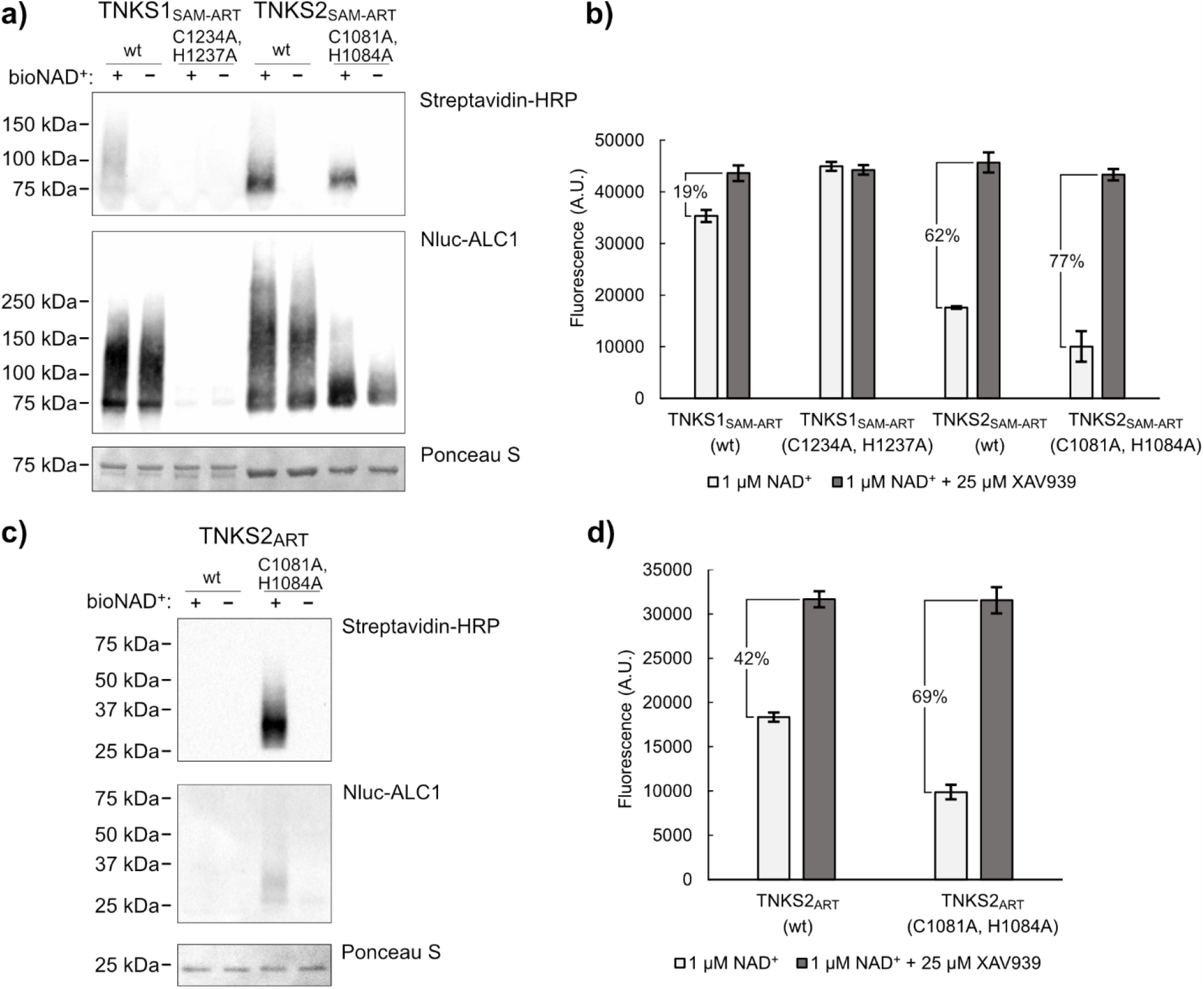
Activity of TNKS1 and TNKS2 constructs. (a) Western blot of TNKS1_SAM-ART_ and TNKS2_SAM-ART_ constructs. TNKS constructs (2 µM) were mixed with 100 nM biotin-NAD^+^ and 900 nM NAD^+^ or buffer as control and incubated for 1 h at room temperature. Auto-modification with biotin-NAD^+^ was detected using streptavidin-HRP. Total modification with PAR was detected using nanoluciferase-ALC1. (b) NAD^+^-consumption assay for TNKS1_SAM-ART_ and TNKS2_SAM-ART_ constructs. The constructs (2 µM) were mixed with 1 µM NAD^+^ and incubated for 1 h at room temperature. The percentage of conversion displayed was determined from the mean fluorescence values. Controls containing 25 µM XAV939 were prepared to determine fluorescence levels without conversion of NAD^+^. (c) Western blot of TNKS2_ART_ constructs. The constructs (2 µM) were mixed with 100 nM biotin-NAD^+^ and 900 nM NAD^+^ or buffer as control and incubated for 20 h at room temperature. (d) NAD^+^-consumption assay for TNKS2_ART_ constructs. The constructs (2 µM) were mixed with 1 µM NAD^+^ and incubated for 20 h at room temperature. The percentage of conversion displayed was determined from the mean fluorescence values. Controls containing 25 µM XAV939 were prepared to determine fluorescence levels without conversion of NAD^+^. Data shown are mean ± standard deviation with number of replicates n = 4. Full-sized blots are shown in **Figure S1**.

In addition to auto-modification, NAD^+^ may also be converted directly by hydrolysis, as several ADP-ribosyltransferases were reported to have NAD^+^ hydrolase activity.^33,34^ To test the total NAD^+^-consumption, PARylation and hydrolysis, we utilized an assay based on chemical conversion of NAD^+^ into a fluorescent compound^35^ (**Figure 2b**). Again, only the mutant of TNKS1_SAM-ART_ did not show any conversion of NAD^+^. Interestingly, despite seemingly lower auto-PARylation activity, the TNKS2_SAM-ART_ mutant showed very similar and possibly higher levels of NAD^+^-consumption compared to the wild type.

As the SAM domain is an important determinant for TNKS activity, we next wanted to investigate the impact of the zinc-binding mutant on the activity without SAM-domain. For this, we similarly to the above tested the auto-PARylation (**Figure 2c**) and total NAD^+^-consumption (**Figure 2d**) for the catalytic domain of TNKS2 and compared wild type and mutant. Due to the lower activity without the SAM domain, a 20-fold longer incubation time was necessary to achieve similar NAD^+^ consumption levels. Strikingly, while auto-PARylation of the wild type catalytic domain could not be detected, a clear smear was present for the zinc-binding mutant. Further, total NAD^+^ consumption was determined to be 69% for the mutant. For the wild type, a lower NAD^+^ consumption of only 42% were measured.

While these findings might not reflect conditions *in vivo*, we believe that these results are indication that activity of TNKS2 is affected upon mutation of the zinc-binding motif. The catalytic domains of TNKS1 and TNKS2 are highly similar, and we have no reason to assume that their molecular mechanism of catalysis would be different. So why do we observe complete abolishment of activity in the TNKS1_SAM-ART_ mutant, while the TNKS2_SAM-ART_ mutant shows comparable or even higher levels of NAD^+^ consumption compared to the wild type? A possible explanation might be improper folding of TNKS1 upon introduction of the mutation. We next set out to test thermal stability of TNKS1 and TNKS2 constructs.

### Thermal stability and folding analysis

To test the thermal stability, we used constructs containing only the ART domain of TNKS1 and TNKS2 both wild type and zinc-binding mutants. We performed differential scanning fluorimetry analysis with all four constructs in absence or presence of TNKS inhibitor XAV939 (**Figure 3a-e**). The melting temperature (T_m_) of TNKS1_ART_ wild type was determined to be 50°C (**Figure 3a**). Analysis of the TNKS1_ART_ mutant however showed an initially high fluorescence value, which decreased as the sample temperature increased with no clear inflection point (**Figure 3b**). As a result, the T_m_ for the TNKS1_ART_ mutant could not be determined. Interestingly, this behaviour in DSF-based experiments is associated with disordered, aggregated or incorrectly folded proteins.^36^ Compared to TNKS1_ART_, the TNKS2_ART_ wild type showed higher thermal stability of 60.1°C (**Figure 3c**), while the zinc-binding mutant showed a dramatic decrease in thermal stability of about 14°C with a T_m_ of 45.8°C (**Figure 3d**). The wild type of TNKS1_ART_ and TNKS2_ART_ as well as the TNKS2_ART_ zinc-binding mutant showed an increased thermal stability of approximately 7-9°C when the potent TNKS inhibitor XAV939 was added, indicating binding. In contrast, the zinc-binding mutant of TNKS1_ART_ did not show changes with XAV939 added. The melting temperatures determined are summarized in **Figure 3e**.

**Figure 3:**
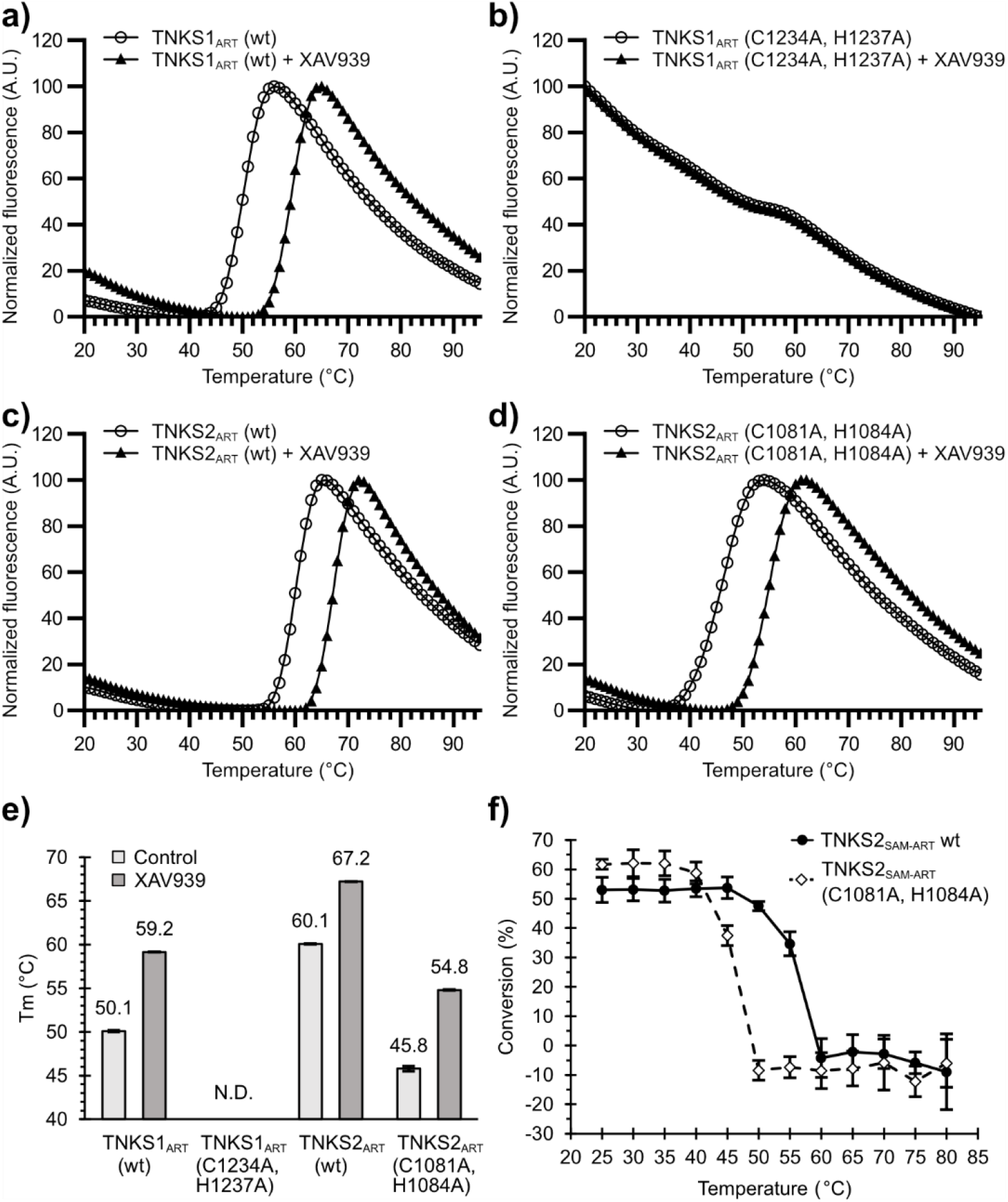
Thermal stability of TNKS1 and TNKS2 ART domain constructs. Representative DSF curves in presence or absence of 25 µM XAV939 for (a) TNKS1_ART_ wild type, (b) TNKS1_ART_ zinc-binding mutant, (c) TNKS2_ART_ wild type and (d) TNKS2_ART_ zinc-binding mutant. (e) Comparison of melting temperatures determined from four independent DSF curves for each condition. Mean melting temperature (T_m_) values are indicated and errors are shown as standard deviations. Melting temperatures for TNKS1_ART_ (f) NAD^+^ consumption assay for TNKS2_SAM-ART_ samples (2 µM) after incubation at different temperatures.

To see if a strong loss of stability in the TNKS2 mutant could also be observed in multimeric TNKS2, we incubated wild type and mutant TNKS2_SAM-ART_ constructs at different temperatures ranging from 20°C to 85°C for 1 min per sample (**Figure 3f**). After incubation, the protein samples were mixed with NAD^+^ and activity was determined using the NAD^+^ fluorescence conversion assay described above. The complete loss of activity is seen for the mutant construct after incubation at 50°C, while the wild type still retained most of its activity after incubation at this temperature. Only after incubation at 60°C did the wild type show complete loss of activity. These results indicate that the zinc-binding motif is a stabilizing structural feature in the case of TNKS2 for both isolated ART domain and multimeric construct containing the SAM and ART domain.

To further characterize possible structural changes introduced by the mutation of the zinc-binding motif in TNKS1 and TNKS2 ART domains, we analysed the TNKS_ART_ constructs using circular dichroism (CD) spectroscopy. Samples were heated from 22°C to 82°C and CD spectra from 190 nm to 260 nm were determined in 2°C intervals. CD spectra at 22°C and 82°C are shown in **Figures 4 a-d**. Spectra for all temperatures are shown in **Figure S2**. The CD spectrum of the TNKS1_ART_ wild type construct (**Figure 4a**) at 22°C shows a minimum at 208 nm, which inverts to a maximum after heating to 82°C. Additionally, minima at 195 nm and 222 nm appear, indicating the thermal unfolding of the protein.^37^ In contrast, the TNKS1_ART_ mutant (**Figure 4b**) at 22°C additionally shows a minimum at 220 nm. After heating to 82°C, no inversion of minima is observed, with only the initial minimum at 208 nm gradually transitioning to 205 nm with no clear inflection point as the temperature increased (**Figure S2**). The spectra of the TNKS2_ART_ wild type (**Figure 4c**) strongly resemble those of the TNKS1_ART_ wild type, with a more pronounced shoulder at 220 nm at 22°C. Interestingly, a major difference in the spectrum is observed at 22°C for the TNKS2_ART_ mutant (**Figure 4d**) compared to the wild type. While no minimum at 208 nm is observed, the construct shows a clear minimum at 220 nm. When heated, the spectrum transitioned comparable to TNKS1_ART_ and TNKS2_ART_ wild type constructs with minima around 195 nm and 222 nm, a maximum at around 210 nm and a clear inflection point (**Figure S2**). To us, these results indicate that mutation of the zinc-binding motif introduces structural changes in the ART domains of TNKS1 and TNKS2. In the case of TNKS1, when considering the DSF result associated with unstructured proteins and only a small gradual change of the CD spectrum upon heating, we believe that mutation of the zinc-binding motif in TNKS1 leads to misfolding of the protein, which likely explains the lack of activity observed in **Figure 2a, b**. Interestingly, the mutation of the zinc-binding motif in TNKS2 leaves the protein active and it is still stabilized by the specific TNKS inhibitor XAV939 in DSF, indicating the presence of a functional binding pocket. Compared to the wild type however, the TNKS2 mutant is severely less thermostable and comparison of the CD spectra at 22°C indicates structural changes, which might explain the different behaviour in catalytic activity observed when compared to the wild type.

**Figure 4:**
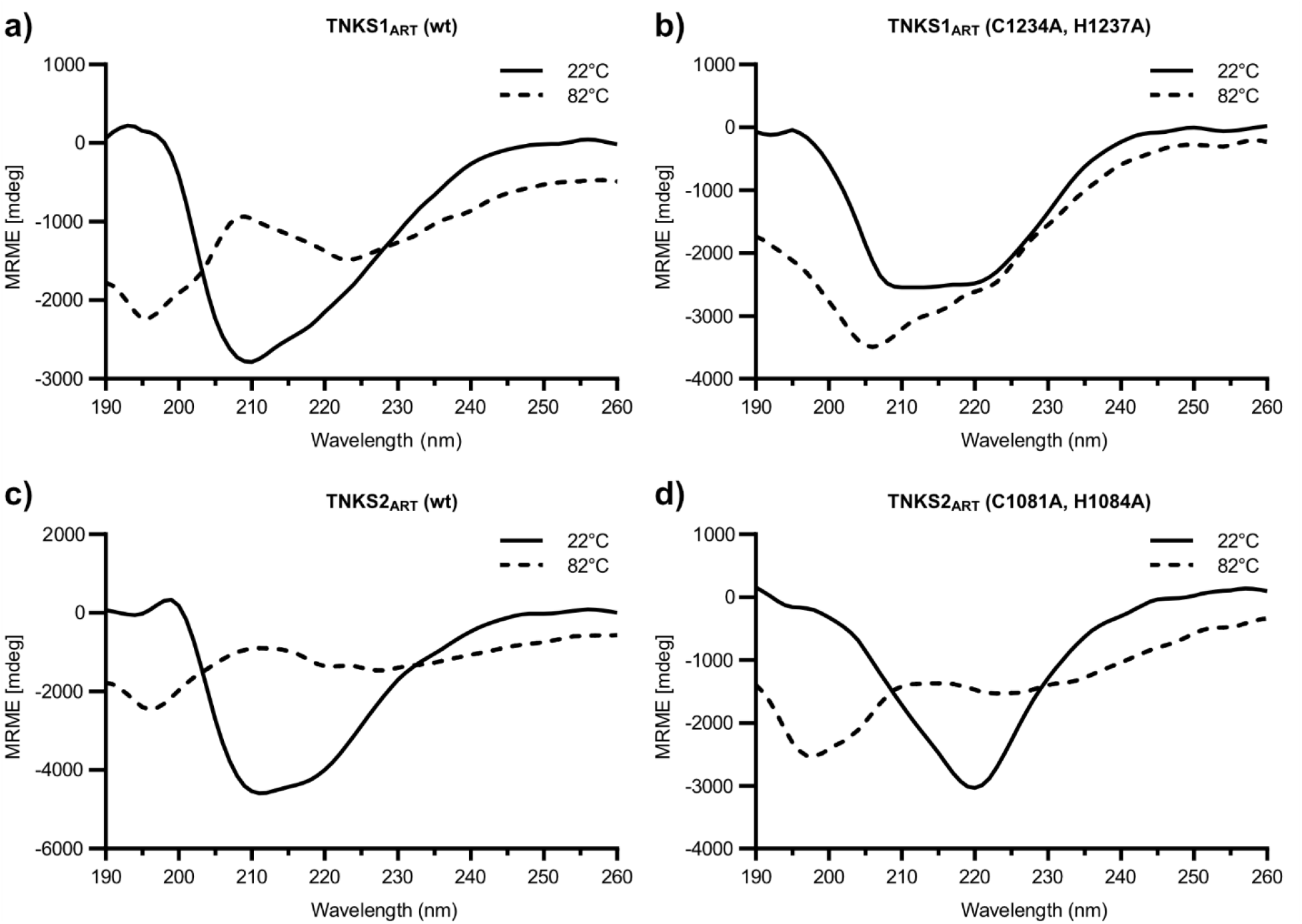
CD spectra of catalytic domain constructs of TNKS1 and TNKS2. The CD spectra for were recorded from 190 nm to 260 nm at 22°C and 82°C for the ART domains of (a) TNKS1_ART_ wild type, (b) TNKS1_ART_ zinc-binding mutant, (c) TNKS2_ART_ wild type and (d) TNKS2_ART_ zinc-binding mutant. CD spectra for all temperatures are shown in **Figure S2**.

### Structural analysis

We aimed to get insight into structural changes occurring in TNKS2 upon disruption of the zinc-binding motif by protein X-ray crystallographic studies. Initial crystallization trials with the zinc-binding mutant of the TNKS2 ART domain remained unsuccessful, which we find not surprising considering the dramatically lower thermal stability compared to the wild type. As alternative strategy, we crystallized the wild type of the TNKS2 ART domain and attempted to remove the zinc through oxidation of the cysteine residues with H_2_O_2_ after protein crystal formation. First attempts with the apo-protein of TNKS2 revealed complete loss of diffraction after H_2_O_2_ treatment, possibly through introduced disorder in the crystal lattice. A different crystal system with TNKS2 can be achieved when co-crystallizing it with inhibitors binding to the adenosine sub-pocket.^38,39^ Treatment with H_2_O_2_ of TNKS2 crystals that were co-crystallized with our previously reported TNKS inhibitor OM-1700 did not significantly lower diffraction quality. While lower concentrations of H_2_O_2_ showed only partial loss of the zinc-ion, incubation of the crystals with 25 mM H_2_O_2_ for 48 hours showed complete loss of electron density corresponding to the zinc ion in the crystal structure. Surprisingly when compared to the structure of non-treated crystals (**Figure 5a**), the structure of H_2_O_2_ treated crystals (**Figure 5b**) showed loss of electron density for the zinc ion (**Figure 5c, d**) and of several protein regions nearby.

**Figure 5:**
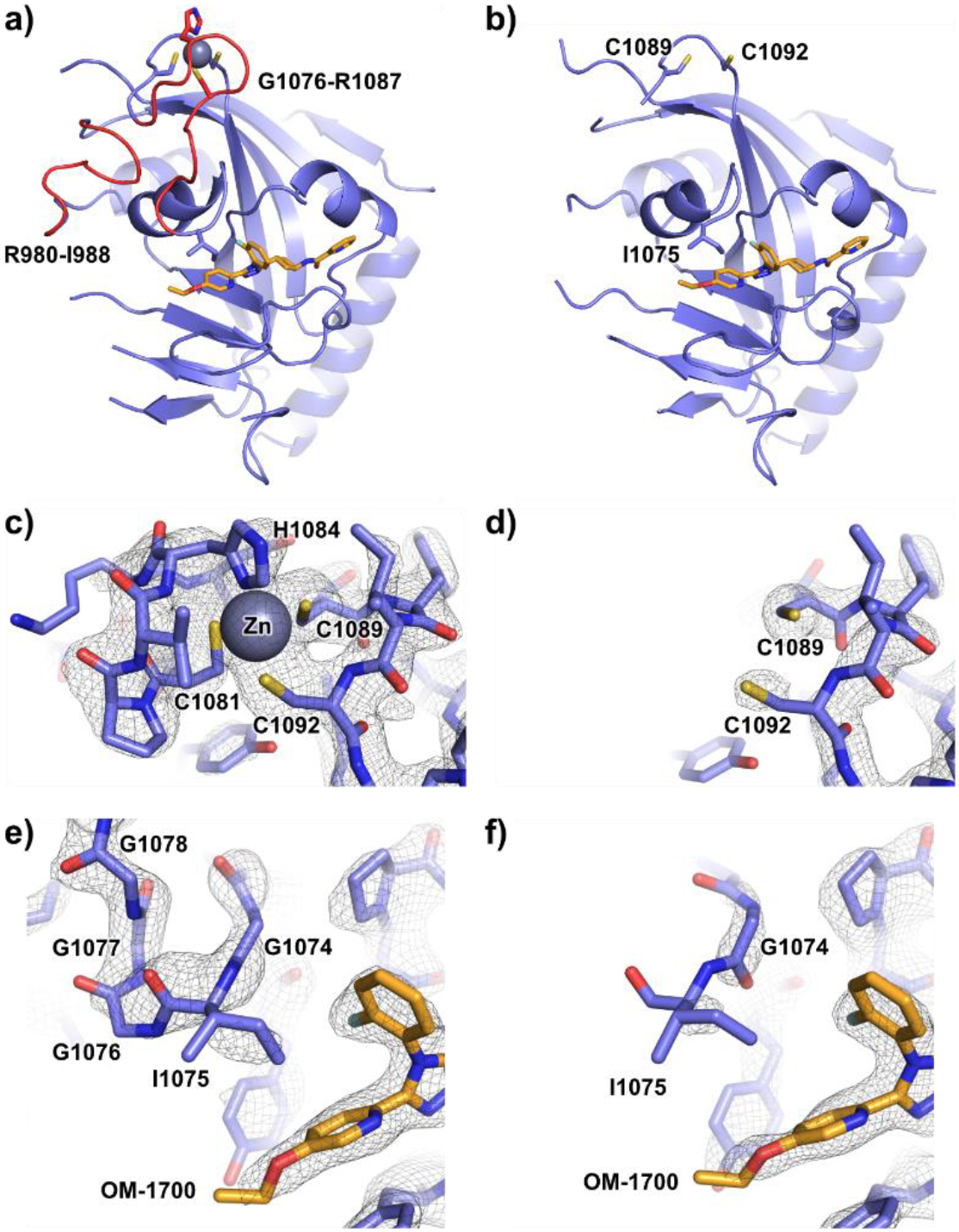
Structural changes in the TNKS2 ART domain upon oxidative removal of zinc. (a) Crystal structure of human TNKS2 ART domain in complex with OM-1700 (PDB ID: 6TG4). Red parts of the protein structure show regions that were not possible to model after H_2_O_2_ treatment. (b) Human TNKS2 ART domain in complex with OM-1700 after the crystals were treated for 48 h with 25 mM H_2_O_2_ (PDB ID: 7POX). The zinc-binding motif region is shown for the structures without (c) and after (d) H_2_O_2_ treatment. A closeup view of the nicotinamide sub-pocket with the bound inhibitor OM-1700 (orange) without (e) and after (f) H_2_O_2_ treatment. The σ_A_ weighted 2F_o_ – F_c_ electron density maps are contoured at 1.6 σ.

Interestingly, residues Gly1076-Arg1087 from the zinc-binding motif up to the Ile1075 located in the G-loop could not be modelled after H_2_O_2_ treatment, indicating increased flexibility of this region (**Figure 5e, f**). Although many of residues in this region are glycines, they form a helix-like structural element with a defined hydrogen bonding network in TNKSs when the zinc-binding motif is intact (**Figure S3**). Additionally, the residues Arg980-Ile988 also showed loss of electron density in chain A of the H_2_O_2_ treated sample (**Figure S4**). This region is located close to the zinc-binding motif and under native conditions is connected by hydrogen bonds to Ser1088 and Arg1087. Electron density for this region was present in chain B, likely due to nearby crystal contacts preventing flexibility.

## Discussion

Based on findings from Kang et al., it was assumed that the zinc-binding motif would be required for the catalytic activity of TNKSs.^30^ Our work shows that the motif is not a strict requirement for the catalytic activity, as PARylation and NAD^+^ consumption were clearly observed for both TNKS2_SAM-ART_ and TNKS2_ART_ mutant constructs. For TNKS1, the activity of the mutant construct was greatly reduced or abolished. Experiments using DSF and CD however showed evidence for incorrectly folded TNKS1 zinc-free mutant protein, whereas the mutant of TNKS2 showed presence of a folded domain, although it had drastically reduced thermal stability. Kang et al. observed complete loss of TNKS1 activity when mutants abolishing the zinc-binding function were used or the zinc was oxidatively removed.^30^ Following these findings, they concluded that the zinc-binding motif in TNKSs may be catalytically essential. Our results however indicate that the TNKS structure, but not directly the catalytic mechanism itself, is affected upon removal of the zinc, which in return may lead to structural defects in TNKS1 causing the loss in catalytic activity.

Analysis of the TNKS2 structure after oxidative removal of the zinc-ion revealed missing election densities for regions Gly1076-Arg1087 and Arg980-Ile988, likely due to increased flexibility in these regions following the disruption of the zinc-binding motif. This loss of structure at the G-loop, which is located at the active site nicotinamide sub-pocket of TNKSs, might affect the activity and can be taken as a possible explanation for the different activities observed in TNKS2 zinc-binding mutant constructs compared to the wild-type proteins.

TNKS ART domains have unique structural adaptations compared to other ARTD family members. These include the zinc-binding motif and the structurally connected G-loop protruding into the active site and forming part of the hydrophobic ‘nook’ in TNKSs. The unique makeup of this region benefitted drug discovery efforts and allowed development of several highly specific TNKS inhibitors.^4,40^ Why do TNKSs have these adaptations compared to other family members? Examining the molecular functions that are unique to the TNKSs might give clues about why these features are present in the TNKS ART domains.

TNKSs and PARP1 and PARP2 are the only ARTD family members in humans that have reported PARylation activity^41^, however TNKSs are evolutionarily more closely related to many of the mono-ADP-ribosylating family members^42^. PARylation activity of TNKSs might have evolved independently from PARP1 and PARP2 and thus might have different underlying structural adaptations for the synthesis of PAR chains. In fact, the PAR chains produced by TNKSs were reported to be linear^5^, while PARP1 and PARP2 are able to produce branched PAR chains^43,44^. This was reasoned to be due to a more restrained acceptor site of TNKSs compared to PARP1 and PARP2, limiting the orientation of a possible acceptor-ADP-ribosyl moiety.^4^ In our western blot experiments, we observed that smears of auto-modified TNKS2_SAM-ART_ zinc-free mutant were running at lower molecular weights compared to the wild type construct, indicating the formation of shorter PAR chains. In PARP1, the acceptor site is required for binding of accepting ADP-ribosyl or PAR moieties during chain elongation.^45^ Assuming the same would be true for TNKSs, an intact zinc-binding region may be required for proper formation of long, unbranched PAR chains.

Another feature of TNKSs is the upregulation of catalytic activity upon multimerization by the SAM domain. It was proposed that a confirmational change in the TNKS ART domains might occur through formation of weak ART domain dimers upon SAM-dependent multimerization^46^, however more work and structural insights are required to confirm this mechanism. We have observed that both mutant and wild-type TNKS2_SAM-ART_ constructs show drastically higher levels NAD^+^ consumption in vitro compared to the TNKS2_ART_ constructs, indicating that the upregulation of activity upon SAM-dependent multimerization is still present when the zinc-binding motif is removed. We note however that the zinc-binding mutant in the TNKS2_ART_ construct showed much higher levels of auto-modification and NAD^+^ consumption compared to the wild type.

Furthermore, the ability of TNKSs to bind plethora of different proteins through their ARC domains and subsequent modification of these proteins is a feature that might require structural adaptations in the makeup of the ART domain. Based on our results, we cannot exclude the possibility that the zinc-binding motif and structurally connected regions play a role in this function or the binding of other yet unknown partners.

In summary, our analysis has revealed that the zinc-binding motif in TNKSs is an important structural feature of the TNKS ART domain and is tightly linked to the structural integrity of the protein and particularly the acceptor site region. Comparing the wild-type and zinc-binding mutant, we observed differences in the PARylation activity of TNKS2, which may be indicative of a role for the zinc-binding motif in catalytic activity regulation, likely through supporting of structural features in the vicinity of the active site. Our findings contribute to the understanding of structural the makeup of the catalytic domains of TNKSs and will help to characterize the basis for TNKSs activity control and regulation in the future.

## Data availability

Atomic coordinates and structure factors have been deposited to the Protein Data Bank under accession number 7POX and raw diffraction images are available at IDA (https://doi.org/10.23729/16748d49-b84c-4114-9431-ecd8ae615623).

## Author contributions

**Sven T. Sowa**: Data curation; formal analysis; investigation; visualization; writing – original draft preparation. **Lari Lehtiö**: Conceptualization; data curation; funding acquisition; supervision; writing – review & editing.

## Acknowledgements

Biocenter Oulu Structural Biology core facility, member of Biocenter Finland, Instruct-ERIC Centre Finland and FINStruct, as well as of “Proteomics and Protein Analysis” and Sequencing core facilities are gratefully acknowledged.

## Material and Methods

### Sequence analysis

Tankyrase sequences were collected from the UniProt database ^47^: *Homo sapiens* TNKS1 (O95271), *Homo sapiens* TNKS2 (Q9H2K2), *Mus musculus* TNKS1 (Q6PFX9), *Mus musculus* TNKS2 (Q3UES3), *Xenopus laevis* TNKS1 (A0A1L8HN28), *Xenopus laevis* TNKS2 (A0A1L8FES1), *Callorhinchus milii* TNKS (A0A4W3JSL5), *Drosophila melanogaster* TNKS (Q9VBP3). The sequences were aligned using Clustal Omega ^48^.

### Cloning

Inserts of the fusion constructs were prepared by PCR: TNKS1_ART_ (1091-1327), TNKS1_SAM-ART_ (1017-1327), TNKS2_ART_ (946-1161) & TNKS2_SAM-ART_ (Met-873-1161). Double mutations of the zinc-binding motif (TNKS1: C1234A, H1237A; TNKS2: C1081A, H1084A) were introduced by overlap-extension PCR using the constructs above as templates. The expression constructs were cloned into pNIC-MBP plasmids using SLIC restriction free cloning method ^49^ as previously described ^19^.

### Protein expression

Expression constructs are based on pNIC-MBP and contain a His_6_-MBP-tag followed by a TEV protease cleavage site (ENLYFQ*SM) before the construct sequence. *E. coli* BL21(DE3) cells were transformed with the plasmids. 500 ml Terrific Broth (TB) autoinduction media including trace elements (Formedium, Hunstanton, Norfolk, England) were supplemented with 8 g/l glycerol and 50 μg/ml kanamycin and inoculated with 5 ml of overnight preculture. The flasks were incubated shaking at 37 °C until an OD_600_ of 1 was reached. The temperature was set to 18 °C and incubation continued overnight. The cells were collected by centrifugation at 4,200×g for 30 min at 4 °C. The pellets were resuspended in lysis buffer (50 mM HEPES pH 7.5, 500 mM NaCl, 15 mM imidazole, 0.5 mM TCEP). Cells were stored at -20 °C until purification.

### Protein purification

All constructs were initially purified by immobilized metal affinity chromatography (IMAC) followed by purification on an MBPTrap column. The cells were thawed and lysed by sonication. The lysate was centrifuged (16,000×g, 4 °C, 30 min), filtered and loaded onto a 5 ml HiTrap HP column equilibrated with lysis buffer and charged with Ni^2+^. The column was washed with 5 column volumes of lysis buffer and 5 column volumes of lysis buffer containing 25 mM imidazole. The protein was eluted using lysis buffer containing 300 mM imidazole. Elutions were directly loaded to a 5 ml MBPTrap HP column. The column was washed with 5 column volumes 20 mM HEPES pH 7.5, 200 mM NaCl, 0.5 mM TCEP and eluted using the same buffer including 10 mM maltose. The elutions of TNKS1_ART_ and TNKS2_ART_ constructs were treated with TEV protease (1:30 molar ratio) for 16-20h at 4 °C. The buffer was exchanged to 20 mM HEPES, 50 mM NaCl, 10% glycerol, 0.5 mM TCEP, pH 7.5 and the protein was loaded to a 5 ml HiTrap SP FF cation exchanger column. The buffer used was 20 mM HEPES, 10% glycerol, 0.5 mM TCEP, pH 7.5 and a linear gradient to a final concentration of 500 mM NaCl over 60 ml was used to elute the protein. To remove residual MBP present, the eluted protein was run over a 5 ml MBPTrap HP column equilibrated with 20 mM HEPES, 500 mM NaCl, 0.5 mM TCEP, 10% glycerol, pH 7.5. Proteins were concentrated, frozen in liquid nitrogen and stored at -70 °C.

### Western blot

For western blot, 10 µl samples were first run in SDS-PAGE (Mini-Protean TGX 4-20% gradient gel, BioRad). The proteins were then transferred to a nitrocellulose membrane using a Mini Trans-Blot Cell Wet/Tank blotting system (BioRad). After transfer, membranes were washed in TBS-T and stained using Ponceau S solution and imaged. Staining was removed by washing the membrane in TBS-T and the membrane was blocked for 20 min using 1% Casein in TBS (BioRad). To detect PAR chains with incorporated 6-Biotin-17-NAD^+^ (Biolog), the membrane was transferred to 15 ml blocking solution including streptavidin-HRP (1:5000) and incubated for 20 min. The membrane was washed using TBS-T buffer, covered with ECL substrate solution (BioRad) and imaged using a ChemiDoc Imaging System (BioRad). The blot was washed with TBS-T and TBS-T including 5% skimmed milk powder. To detect poly-ADP-ribosyl groups, the membrane was transferred to 15 ml TBS-T including 1% skimmed milk powder and nanoluciferase-ALC1 (0.1 µg/ml) ^32^ and incubated for 20 min. The membrane was washed with TBS-T and imaged using 500 µl of 1:500 NanoGlo substrate (Promega, catalogue number: N1120) diluted in 10 mM sodium phosphate buffer pH 7.0.

### NAD^+^ conversion assay

The NAD^+^ conversion assay is based on our previously described method ^35^. Briefly, 5 µl reactions were prepared in 384-well ShallowWell black polypropylene plates (Fisherbrand). TNKS_SAM-ART_ or TNKS_ART_ constructs (2 µM) were mixed with 1 µM NAD^+^ and let incubate at room temperature for 1 h or 20 h, respectively. Controls containing 25 µM XAV939 preventing the NAD^+^ conversion were prepared and used for the calculation of the NAD^+^ conversion percentage. The reactions were stopped by addition of 1 µl KOH (2 M) and 1 µl ethanol containing 10%(v/v) acetophenone and 30%(v/v) glycerol. The mixtures were incubated for 10 min at room temperature. Finally, 3 µl formic acid (100%) were added and the reactions incubated for 5 min at room temperature. The fluorescence was read using a Spark multimode plate reader (Tecan) with excitation wavelength of 372 nm (10 nm bandwidth) and emission wavelength of 444 nm (20 nm bandwidth). The reactions were prepared using a Mantis microfluidic liquid handler (Formulatrix).

### Differential scanning fluorimetry

All samples of TNKS1_ART_ or TNKS2_ART_ constructs were prepared in 20 mM HEPES pH 7.5, 500 mM NaCl, 0.5 mM TCEP, 10% glycerol. 5 µM of the constructs were mixed with SYPRO orange (2x) in presence of absence of 25 µM XAV939. Samples were transferred to 96-well qPCR plates. Measurements were performed in a BioRad C1000 CFX96 thermal cycler. Data points for melting curves were recorded in 1 min intervals from 20–95 °C, with the temperature increasing by 1 °C/min. The analysis of the data was done in GraphPad Prism 7 using a nonlinear regression analysis (Boltzmann sigmoid equation) of normalized data.

### Circular dichroism spectroscopy

TNKS1_ART_ or TNKS2_ART_ constructs were diluted to 20 µg/ml in 10 mM sodium phosphate buffer (pH 7.5). CD spectra were recorded using a Chirascan™ CD Spectrometer (Applied Photophysics Ltd). Spectra (190 nm to 260 nm) were recorded in 2 °C intervals from 22–82 °C, with the temperature increasing by 1 °C/min.

### Crystallography and data collection

Protein crystallization for TNKS2_ART_ in complex with OM-1700 was done as previously described ^39^. Protein crystals were soaked in reservoir solution containing 25 mM H_2_O_2_ for 48 hours. Data were collected at Diamond Light Source on beamline i03. Diffraction data were processed using the XDS package ^50^. All structures were solved using molecular replacement with PHASER ^51^ using the structure of TNKS2 (PDB code: 5NOB) as a starting model. Coot ^52^ and Refmac5 ^53^ were used for model building and refinement, respectively. The images of the structures were prepared using PyMOL (The PyMOL Molecular Graphics System, version 1.8.4.0, Schrödinger, LLC.). Data collection and refinement statistics are shown in **Table S1**.

