## Supplementary information for "The zinc-binding motif in tankyrases is required for the structural integrity and proper function of the catalytic domain"

### Content

**Figure S1: Full sized blots.**

**Figure S2: CD spectra of catalytic domain constructs of TNKS1 and TNKS2.**

**Figure S3: The glycine-rich region adjacent to the zinc-binding motif in tankyrases is structured under native conditions.**

**Figure S4: Loss of electron density in the TNKS2 ART domain region Arg980-Ile988 upon oxidative removal of zinc.**

**Table S1: Data collection and refinement statistics.**

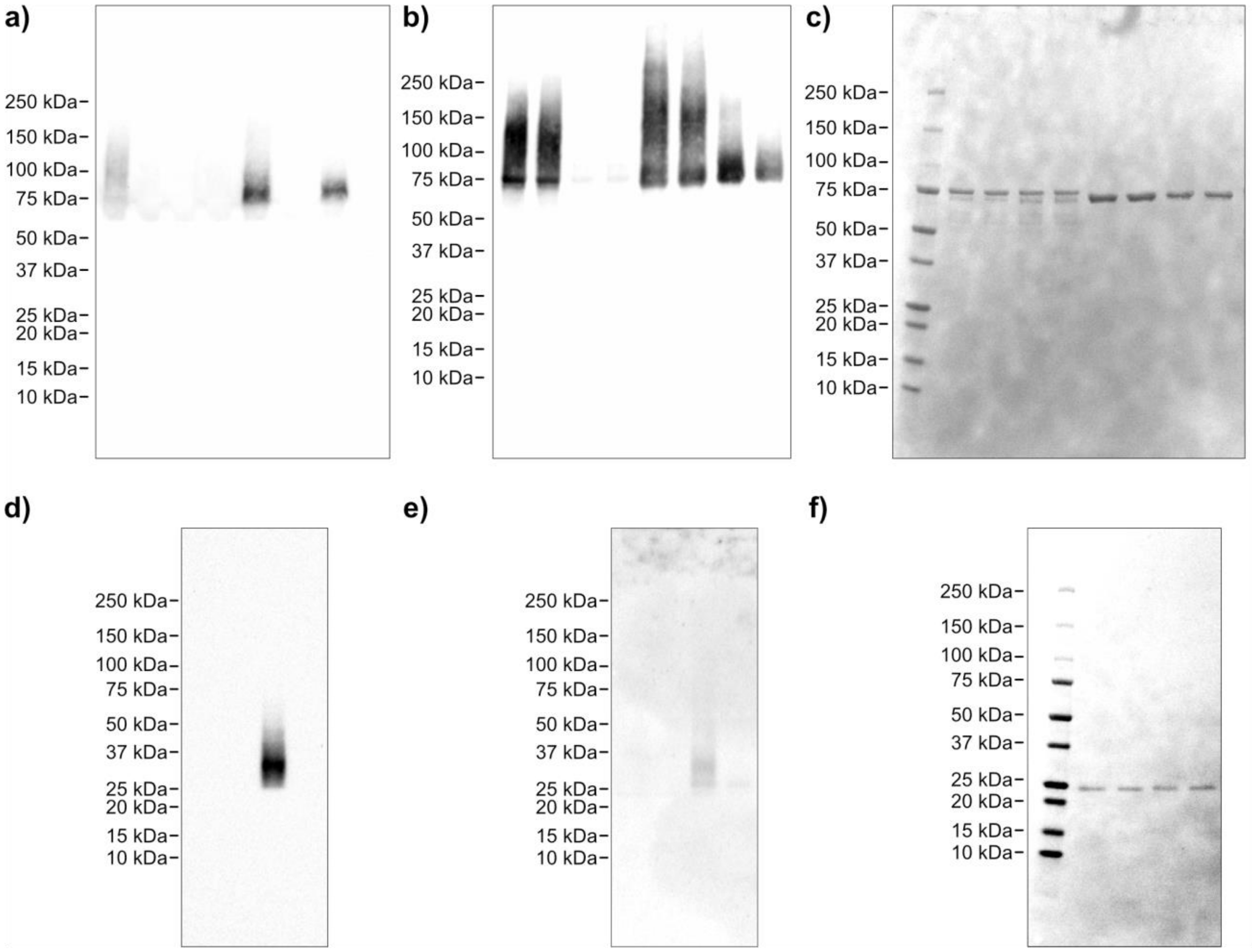


**Figure S1: Full-sized blots.** Full-sized blots related to **Figure 2a**: (a) Detection with streptavidin-HRP. (b) Detection with Nluc-ALC1. (c) Ponceau S staining. Full sized blots related to **Figure 2c**: (d) Detection with streptavidin-HRP. (e) Detection with Nluc-ALC1. (f) Ponceau S staining.

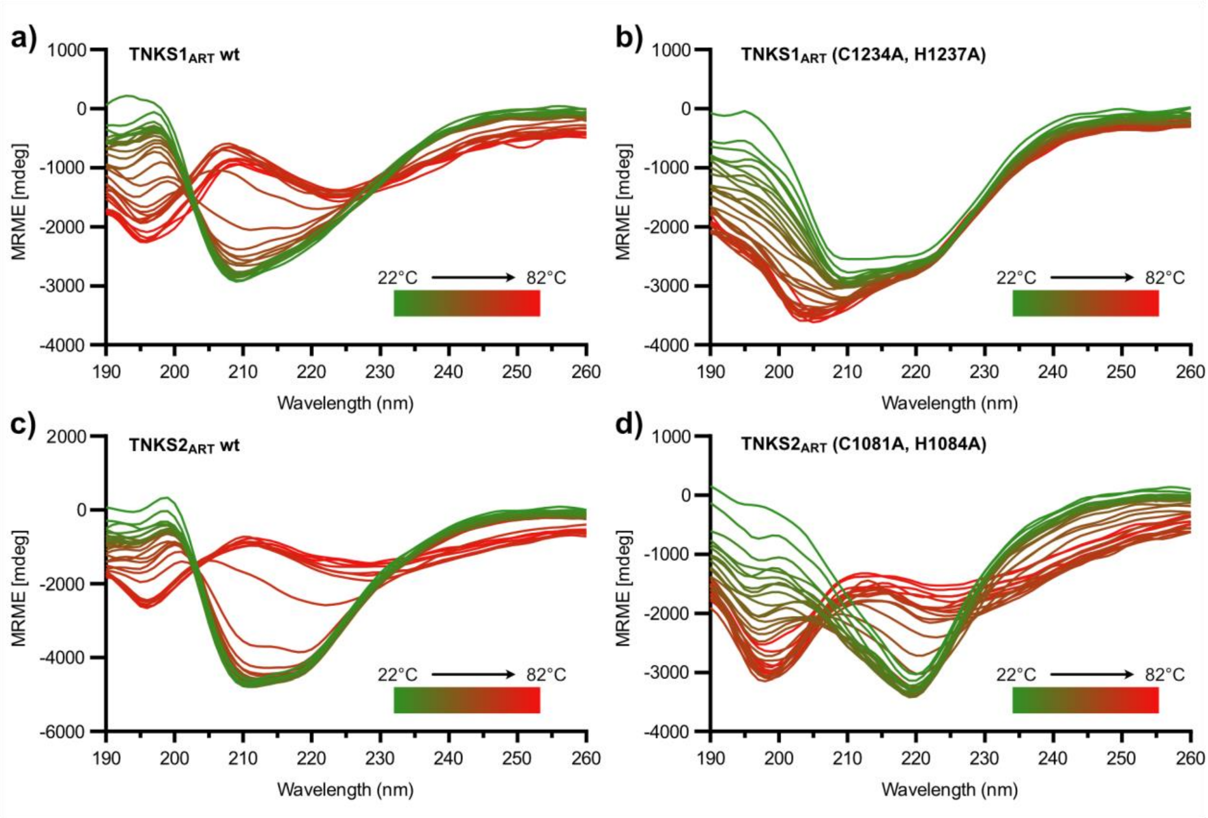


**Figure S2: CD spectra of catalytic domain constructs of TNKS1 and TNKS2.** The CD spectra for were recorded from 190 nm to 260 nm at 22°C in increments of 2°C up to 82°C for the ART domains of (a) TNKS1 wild type, (b) TNKS1 zinc-binding mutant, (c) TNKS2 wild type and (d) TNKS2 zinc-binding mutant.

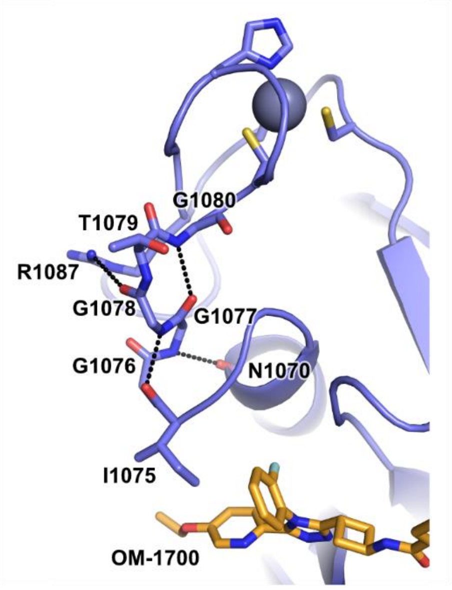


**Figure S3: The glycine-rich region adjacent to the zinc-binding motif in tankyrases is structured under native conditions.** The glycine-rich region (TNKS2: G1076-G1080) forms a helical structure with hydrogen bond network in tankyrases under native conditions. After removal of the zinc-ion by $H_2O_2$ treatment, the electron density is not present for this region. The crystal structure shown is of the human TNKS2 ART domain in complex with OM-1700 (PDB ID: 6TG4). Hydrogen bonds are shown as black dashes.

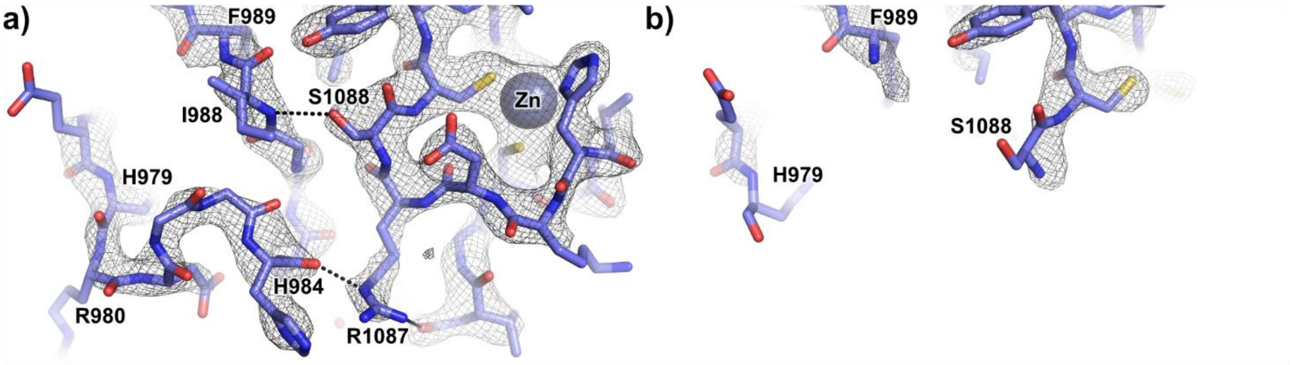


**Figure S4: Loss of electron density in the TNKS2 ART domain region Arg980-Ile988 upon oxidative removal of zinc.** (a) Crystal structure of human TNKS2 ART domain in complex with OM-1700 (PDB ID: 6TG4). (b) Human TNKS2 ART domain chain A in complex with OM-1700 after the crystals were treated for 48 h with 25 mM H2O2 (PDB ID: 7POX). Hydrogen bonds are shown as black dashes. The $\sigma_A$ weighted $2F_o - F_c$ electron density maps are contoured at 1.6 σ.

**Table S1: Data collection and refinement statistics.**

| | TNKS2 (H2O2 treated)<br>PDB ID: 7POX |
|---|---|
| **Data collection** | |
| Resolution (Å) | 41.41 – 2.5 |
| (Outer shell) | (2.589 – 2.5) |
| Wavelenght | 0.97625 |
| Beamline | DLS, i03 |
| Temperature (K) | 100 |
| Space group | P $2_12_12_1$ |
| Cell dimensions | |
| *a*, *b*, *c* (Å) | 41.64, 76.44, 147.79 |
| α, β, γ (°) | 90, 90, 90 |
| No. of unique reflections | 17031 (1670) |
| $R_{merge}$ | 0.200 (1.467) |
| Mean $I/\sigma I$ | 10.77 (1.91) |
| Completeness (%) | 98.92 (98.75) |
| Redundancy | 13.3 (13.8) |
| $CC_{1/2}$ (%) | 99.8 (90.0) |
| Wilson B-factor (Å$^2$) | 48.28 |
| **Refinement** | |
| $R_{work}$ / $R_{free}$ | 0.247 / 0.294 |
| No. non-H atoms | 3210 |
| protein | 3110 |
| ligands/ions | 68 |
| waters | 32 |
| *B*-factors (Å$^2$) | |
| protein | 60.10 |
| ligands/ions | 51.61 |
| waters | 45.82 |
| RMSD bonds / angles | 0.013 / 1.61 |
| Ramachandran plot (%) | |
| favored/allowed/outliers | 96.9 / 3.1 / 0.0 |

*Values in parentheses are for the highest-resolution shell.
